# Towards explaining the fertility gap in farmed Pekin ducks

**DOI:** 10.1101/2024.09.12.612586

**Authors:** Cammy H Beyts, Jonathan Wright, Yimen Araya-Ajoy, Kellie Watson

## Abstract

Maximising reproductive success is crucial to animal production systems, particularly in meeting global demands for animal products and improving commercially important traits. However, while social interactions and mating strategies are known to influence reproductive success in wild populations, their consideration in agricultural systems remains limited. Using an interdisciplinary framework that combines concepts from behavioural ecology and quantitative genetics in an animal breeding context, we investigated the role of sperm limitation and polygynous mating strategies (female polyandry, male monopolisation of females and male polygamy) in limiting female reproductive success in farmed Pekin ducks (*Anas platyrthynchos domestica*). We assessed the impact of these behaviours on chick production and quantified their genetic and environmental (co)variance. Our results revealed that the number of dam mates positively influenced chick production in female ducks. However, contrary to our expectation, skew in chick paternity (our measure of male monopolisation) was associated with increased female chick production, challenging the hypothesis that male monopolisation limits the sperm available to females and reduces their reproductive success. We found no evidence that male polygamy led to decreased female chick production. Genetic analysis revealed that female mate number and reproductive skew exhibit genetic variance, providing opportunities for targeted selection to enhance chick production. However, there was a negative genetic association between female polyandry and skew in chick paternity, suggesting a trade-off between these traits that would need to be considered in future selection programmes. Our findings highlight how concepts from behavioural ecology can be incorporated into breeding programmes, providing new opportunities to develop effective and sustainable breeding strategies.

## Introduction

Maximising reproductive success is vital to animal production systems, enabling breeders to meet global demand for animal products and facilitating improvements in commercially important production traits such as growth rate, food conversion efficiency and meat yield, whilst simultaneously ensuring that animal welfare remains high (Rauw et al. 1998; Diskin and Morris 2008; Davis and White 2020; Fernandez-Novo et al. 2020). In behavioural ecology, complex behavioural traits that influence social interactions and patterns of mating are vital for understanding factors affecting reproductive success in natural populations (Sih et al. 2009; Alberts 2019; Philson and Blumstein 2023). Researchers can assess the fitness consequences of these social interactions, such as dominance hierarchies and mating strategies, to determine their adaptive significance and predict responses to natural selection (Sinervo and Zamudio 2001; Westneat and Stewart 2003; Kokko and Jennions 2008; Chen et al. 2011). By employing quantitative genetic methods like pedigree-based analyses alongside individual-level phenotypic data, it is possible to gain further insight into the genetic basis of these behaviours and their potential to evolve under different selection pressures (Falconer and Mackay 1996; Hill 2010; Wilson et al. 2010). However, despite the widespread use of quantitative genetics in animal breeding, social interactions are rarely considered in agricultural systems (Turner et al. 2009; Camerlink et al. 2015; Angarita et al. 2019; Canario et al. 2020). This is surprising given that many breeding programmes rely on natural matings between group-housed animals, where social interactions are likely to be central in determining individual reproductive success (Brillard 2003; Brillard 2004; Bilcik et al. 2005). By considering the adaptive significance of social interactions, breeders can gain more accurate understanding of reproductive success, welfare outcomes and economically important traits, ultimately increasing the efficiency and sustainability of the industry (Rauw et al. 1998; Knap 2005; Reijrink et al. 2009).

Pekin ducks (*Anas platyrhynchos domesticus*) reared for meat and egg production are a commercially important species (Makagon and Riber 2022). Despite their prolific egg-laying capabilities, a considerable proportion of these eggs remain unfertilised, resulting in reduced hatchability rates and productivity, which is frequently referred to as the “fertility gap” (Abd El-Hack et al. 2019). With a strong and growing demand for day-old ducklings in the industry, there is increasing interest in identifying strategies to improve fertility and increase viable chick numbers (Jones and Dawkins 2010; Campbell et al. 2015; Abd El-Hack et al. 2019).

The low fertilisation success in Pekin ducks may be influenced sperm limitation in females (Cartar 1985; Moller 1991; Preston et al. 2001). The sperm limitation hypothesis suggests that female reproductive output is constrained by sperm availability, potentially due to females receiving insufficient sperm during a single mating to fertilise all their eggs. As such, realised female reproductive success may be lower than its potential (Wedell et al. 2002; Levitan 2004; Parker and Pizzari 2010). To counteract sperm limitation, females may engage in polyandry, mating with multiple males to increase the chance of acquiring enough sperm to fertilise all their eggs (Wedell et al. 2002; Alonzo and Pizzari 2013; Bocedi and Reid 2016). Consequently, a positive association between the number of mating partners a female Pekin duck has and the number of offspring she produces might be expected.

In response to female polyandry, males may attempt to monopolise females to limit female access to rival males (Arnqvist and Rowe 1995; Rowe et al. 2005; Engelhardt et al. 2006; Kvarnemo and Simmons 2013; Lüpold et al. 2014). This may increase a male’s chances of paternity by avoiding costly postcopulatory sperm competition (Parker 2020; Dougherty et al. 2022). However, male monopolisation could negatively impact female reproductive success if the monopolising male fails to provide enough sperm to fertilise all a female’s eggs (Kvarnemo and Simmons 2013). Consequently, increased female monopolisation by males, as indicated by males which tend to dominate and sire a greater proportion of chicks in a dam’s brood (i.e., higher skew in chick paternity), might be expected to be associated with a decrease in the number of chicks a female produces.

Constraints on sperm production may also result in male sperm counts being depleted as a result of frequent or successive copulations (Dewsbury 1982; Birkhead and Fletcher 1995; Preston et al. 2001). Thus, when males have lots of mating partners, the amount of sperm they can transfer to each female or mating event may decrease, exacerbating sperm limitation in females (Preston et al. 2001; Wedell et al. 2002). Consequently, we might also expect that females that mate with males, who themselves have a high number of mating partners, will have fewer chicks compared to females who mate with males with fewer mating partners.

Understanding how to bridge the fertility gap in farmed Pekin ducks therefore requires consideration of how female mating strategies (e.g. polyandry) may maximise sperm availability, as well as how male mating strategies (e.g. mate monopolisation and male polygamy) may act as constraints on female reproductive success (Wedell et al. 2002; Kvarnemo and Simmons 2013). Investigating the contribution of sperm limitation on female reproductive success through the interplay between polyandry, male monopolisation and polygamy should offer valuable insights for enhancing fertility and chick production in farmed Pekin ducks.

Ultimately resolving the fertility gap requires quantifying the genetic (co)variance associated with the different mating behaviours because this determines the potential of the mating system to respond to selection (Merilä and Sheldon 2000; Charmantier and Garant 2005; McAdam et al. 2011). This will in turn inform management practices and influence breeding system design (Rydhmer 2000; Safari et al. 2005; Berry et al. 2014). For example, if some mating strategies, such as female mate number, are associated with higher female reproductive success, breeding programmes can identify and select for individuals with a genetic predisposition for these traits.

Furthermore, by understanding the genetic correlations between mating strategies, breeders can also establish whether genetic progress on both traits can be targeted simultaneously or whether improvements in one trait may come at the expense of the other (Falconer and Mackay 1996; Lynch and Walsh 1998; Walsh 2008). For example, assuming that polyandry is associated with an increase in female chick production, but male monopolisation is associated with a decrease in chick production, a positive genetic correlation between the number of mates females have and skew in chick paternity would mean that selection for females that show high levels of polyandry would also lead to selection for females that are vulnerable to high levels of male monopolisation. Thus, genetic progress in chick production may be limited if selection for females with high numbers of mating partners simultaneously selects for high rates of male monopolisation. Understanding how traits are genetically correlated therefore allows breeders to make more informed decisions on how to optimise breeding programmes (Falconer and Mackay 1996; Walsh 2008; Hill 2010).

The aim of this study is to assess how much sperm limitation contributes to the fertility gap in farmed Pekin ducks and determine the evolutionary potential to solve this problem. We therefore have three key objectives: i) examining the influence of the number of mating partners on female chick production; ii) examining the effect of skewness in chick paternity (as a measure of female monopolisation by males) on female chick production; and iii) examine how male polyandry impacts female chick production. From the sperm limitation hypothesis, we predict that an increase in female polyandry will correspond to an increase in chick production and that greater skew in chick paternity will be associated with a decline in chick production. We further predict that females that mate with males who have lots of mating partners themselves will suffer a further decline in chick production. In addition to these primary objectives, we use pedigree data to explore the extent to which female mate number, skew in chick paternity and male mate number has a genetic (heritable) basis, as well as whether there are genetic covariances between these effects on female chick production. Exploring the genetic contribution of these traits, as well as their genetic covariances, has the potential to help guide selection strategies and management practices enhancing the productivity, welfare and reproductive success of farmed Pekin ducks.

## Methods

### Study population

We conducted our study on a UK population of commercially farmed pedigree Pekin ducks. Data collection spanned from 2019 and 2022 and included five generations of ducks across four commercial Cherry Valley pedigree Pekin duck strains. The ducks were housed in group pens (n = 36) with each pen housing between 124 and 291 adult individuals (mean = 155.6, SD = 38.02) at a ratio of 4:1 females to males. The birds were group housed over a 12-week period during which time their movements were unrestricted within the pen, allowing for natural social interactions and mating behaviours. At the beginning of the study period, birds ranged in age from 28 to 38 weeks. Food access was timed and strain dependent, provided via a feed hopper within the pen, while water was available *ad libitum* through suspended nipple drinkers.

During the 12-week period, all eggs laid were collected daily and every 10-14 days were transferred to a setter incubator for approximately 25 days. At day 10, eggs were candled to remove infertile (clear) and infected eggs. After the setter stage, eggs were moved to the hatcher where they stayed 3-4 days until hatching. Any remaining infected eggs were removed on day 35. Infertile and infected eggs were not included in the final data set.

### Pedigree reconstruction and deriving reproductive and behavioural parameters

Blood samples were collected from adult males, females and day-old chicks in the studied population. Low-density SNP arrays were used for genotyping the samples. This genomic data was used to reconstruct the pedigree, enabling the determination of the dam and sire of each chick as well as the genetic relatedness between individuals. Genotyping was performed using a low-density SNP panel consisting of 450 SNPs. Blood samples were obtained from the leg of each duck and duckling using sterile lancets (28G) and deposited on Qiagen FTA cards cut to a size of 5×10mm (Non-indicating FTA Cards, qiagen.com). The FTA cards were then placed in a 96-well plate and sent to ThermoFisher for Eureka genotyping services (Eureka^TM^ Genotyping services, Catalog Panels, Themofisher.com). This genomic data was used to reconstruct the pedigree to determine the dam and sire of each chick as well as assess the genetic relatedness between individuals using the Sequoia package (Huisman 2017) in RStudio.

Using the reconstructed pedigree, we derived several parameters to determine an individual’s reproductive success and mating behaviour: i) the total number of chicks produced by each dam as a measure of female reproductive success; ii) the number of mating partners that resulted in viable offspring for each dam as measure of female polyandry; iii) the number of mating partners that resulted in viable offspring for each sire as measure of male polygamy; and iv) the coefficient of variation in chick paternity per dam as a measure of reproductive skew or male monopolisation per female. To calculate the coefficient of variation in chick paternity, we first calculated the standard deviation in the number of chicks sired by each male a female mated with. We then divided this standard deviation by the mean number of chicks females produced across all her mating partners. This measure of reproductive skew allowed us to quantify the relative variation in chick paternity across males for each female. We chose the coefficient of variation in chick paternity as our measure of reproductive skew as this provided us with an assessment of skew in male paternity that was independent of the total number of mates a female had, enabling us to better understand the distinct contributions of female polyandry and male monopolisation on female reproductive success (Vehrencamp 1983; Pamilo and Crozier 1996).

## Statistical analysis

### Mating behaviours influencing chick production in dams

To investigate the effect of different mating behaviours on chick production, whilst also considering the relative contributions of genetic and environmental factors, we used a series of animal models. An animal model is a type of mixed model that allows incorporating pedigree information to estimate additive genetic variation and covariation between traits (Wilson et al. 2010). In each model, we used the total number of chicks produced by each dam as the response variable, which was treated as count data and fitted with a Poisson distribution. To account for genetic variability among females in their ability to produce chicks, we included a random effect of Dam ID, linked to the pedigree. Additionally, we included a random effect of Sire ID, linked to the pedigree, to assess whether there was genetic variation among males in their impact on female reproductive success. To capture the different social interactions that may be occurring within specific pens, we included a random effect of Pen ID in the models to capture this potential source of environmental variance.

A series of these animal models with increasingly complex fixed effect structures was used to test for possible effects of sperm limitation (see Introduction). In the base model (Model 1) we used an intercept only model, which quantified the relative contribution of Dam ID, Sire ID and Pen ID on dam chick production. In Model 2, we included the total number of dam mating partners as a fixed effect to determine how female polyandry contributed to dam chick production. In Model 3, we extended the model to include the coefficient of variation in chick paternity as a fixed effect as a measure of skew in chick paternity or male monopolisation. In Model 4, we looked at how female chick production was influenced by male sperm depletion by looking at the effect number of mates their mating partners had on female reproductive success. For this behavioural parameter, we chose to focus on the number of mating partners of a female’s dominant mate. This approach allowed us to assess the potential influence of a particular male’s behaviour, as opposed to the collective impact of multiple mating partners on female reproductive success. Finally, as we found a significant effect of female polyandry and skew in male paternity on female reproductive success (see results section), we also included Model 5 that included a fixed effect interaction term between the total number of dam mates and the coefficient of variation in chick paternity.

### Genetic variance and covariance between mating behaviours

To understand the genetic and environmental factors influencing the total number of dam mates, skew in chick paternity and the number of mating partners of the female’s dominant male, we conducted univariate intercept-only models for each of these traits. To estimate the genetic variance associated with each mating behaviour, we fitted models with Dam ID as a random effect, linked to the pedigree, to assess the genetic variation of these mating behaviours among females. We also fitted Pen ID as a random effect to capture the environmental variation resulting from social interactions within pens. When examining the genetic variance associated with the number of mating partners of the female’s dominant male, we also included a random effect of Sire ID linked to the pedigree. This was because the number of sire mates was a predominantly male trait and allowed us to determine the genetic predisposition for males to attract and mate with multiple females.

The results of the univariate analysis guided the design of our subsequent bivariate analysis, which aimed to explore the genetic and phenotypic covariate between two key traits identified in the univariate analysis, female polyandry and skew in chick paternity. To achieve this, we utilised a bivariate animal model, incorporating the number of dam mates and coefficient of variation in variation in chick paternity as response variables (Thompson et al. 1995; Roman and Wilcox 2000; Wilson et al. 2010). To estimate the genetic covariance between these traits at the female and male level, we fitted a random effect of Dam ID and Sire ID linked through the pedigree. Furthermore, we also looked at the environmental variance between these traits at the Pen level by incorporating a random effect of Pen ID.

### Mating behaviours influencing chick production in sires

As our findings showed mixed support for the sperm limitation hypothesis (see Results section), we explored additional hypotheses by examining the impact of the same mating behaviours on male reproductive success by using the same 5 animal models detailed above, but with reciprocal variables based upon the males rather than the females. In each model, we used the total number of chicks sired by each male as the response variable and used random effects of Sire ID and Dam ID, linked through the pedigree, as well as Pen ID to estimate genetic and environmental sources of variation in male chick production.

Similar to the models examining female reproductive success, we used an intercept only as well as three additional models with fixed effects to examine how the number of sire mates, skew in chick maternity (measured as the coefficient of variation in chick maternity) and female polyandry (measured as the average number of mating partners of the male’s female mates) impacted male reproductive success. We also examined how the interaction between the number of sire mates and skew in chick maternity impacted the number of chicks males sired.

We anticipated that higher number of sire mates would obviously lead to an increase in male reproductive success. However, if monopolisation of females is a mating strategy used to reduce sperm competition from rival males, then we would expect that males that sire a higher proportion of a dam’s brood will have higher reproductive success. Similarly, we might also expect that males that mate with females who they themselves have lots of mates would be exposed to higher levels sperm competition, reducing male reproductive success.

Finally, we also examined genetic and environmental variance in sire mating behaviours, as well as the genetic and environmental covariance between the number of sire mates and skew in chick paternity in a reciprocal approach already described for females.

## Results

### Mating behaviours influencing chick production in dams

In the base model (Model 1, Table 1), we found non-zero genetic variance associated with Dam ID. This indicates genetic differences among dams in their capacity to produce chicks. There was also genetic variance associated with Sire ID, suggesting that certain sires can positively influence the reproductive success of their mates, contributing to increased chick production. However, the genetic effect of Sire ID was an order of magnitude less than that observed in Dam ID. We also found an environmental source of variance in chick production among pens, which may indicate that the particular social interactions between birds within each pen may contribute to female chick production.

**Table 1.** Results of five animal models of increasing complexity examining the impact of different mating behav iours (fixed effects) and sources of genetic and environmental variation (random effects) on dam chick production in farmed Pekin ducks. Model number corresponds to the model used in each analysis.95% c redible intervals are reported in parentheses.

| Model | 1 | 2 | 3 | 4 | 5 |
| --- | --- | --- | --- | --- | --- |
| **Fixed effects** |  |  |  |  |  |
| Intercept | 2.840 (2.72, 2.97) | 3.264 (3.18, 3.36) | 2.612 (2.54, 2.68) | 2.616 (2.55, 2.700) | 2.635 (2.57, 2.71) |
| Number of dam mates |  | 0.144 (0.14, 0.15) | 0.122 (0.12, 0.13) | 0.122 (0.12, 0.13) | 0.149 (0.14, 0.16) |
| Skew in chick paternity |  |  | 0.842 (0.81, 0.88) | 0.843 (0.81, 0.88) | 0.629 (0.59, 0.67) |
| Number of mates of most dominant sire |  |  |  | 0.0005 (-0.0002, 0.001) | 0.00623 (-0.00004, 0.001) |
| Number of dam chicks : skew in chick paternity |  |  |  |  | -0.0527 (-0.06, -0.04) |
| **Random effects** |  |  |  |  |  |
| Pen ID | 0.25 (0.238, 0.264) | 0.069 (0.04, 0.11) | 0.038 (0.02, 0.06) | 0.039 (0.02, 0.07) | 0.04 (0.02, 0.06) |
| Dam ID $V_G$ | 0.166 (0.14, 0.20) | 0.054 (0.05, 0.06) | 0.019 (0.01, 0.02) | 0.016 (0.01, 0.02) | 0.014 (0.01, 0.02) |
| Sire ID $V_G$ | 0.017 (0.02, 0.02) | 0.007 (0.0001, 0.01) | 0.004 (0.002, 0.007) | 0.005 (0.003, 0.008) | 0.005 (0.002, 0.007) |
| Residual | 0.231 (0.20, 0.26) | 0.104 (0.09, 0.12) | 0.071 (0.06, 0.08) | 0.074 (0.07, 0.08) | 0.05 (0.05, 0.06) |
| **Quantitative parameters** |  |  |  |  |  |
| Repeatability: Pen ID | 0.25 (0.238, 0.264) | 0.252 (0.246, 0.260) | 0.214 (0.185, 0.241) | 0.214 (0.187, 0.240) | 0.205 (0.177, 0.23) |
| Heritability: Dam ID | 0.256 (0.247, 0.266) | 0.249 (0.245, 0.252) | 0.259 (0.244, 0.275) | 0.269 (0.255, 0.282) | 0.269 (0.257, 0.284) |
| Heritability: Sire ID | 0.22 (0.216, 0.225) | 0.237 (0.235, 0.240) | 0.355 (0.329, 0.392) | 0.347 (0.327, 0.369) | 0.342 (0.317, 0.368) |

In Model 2 (Table 1), females that had more mates also tended to have more chicks. This highlights the adaptive benefits of polyandry on reproductive success in farmed female Pekin ducks. Furthermore, we found that inclusion of the number of mates as a fixed effect reduced the genetic variances associated with Dam ID and Sire ID, as well as the variance associated with Pen ID in comparison to the base Model 1 (Table 1). This emphasises the role of female mate number in explaining female reproductive success, as well as the genetic and environmental variance associated with chick production.

In Model 3 (Table 1), we found that higher levels of reproductive skew were positivity associated with an increase in chick production. This suggests that when one or a few males sired the majority of a female’s chicks, females had higher reproductive success than when paternity of her chicks was distributed more evenly among males. Notably, this result partly contradicts our initial prediction from the sperm limitation hypothesis, where we expected a negative association between skew in male paternity (potentially indicative of male monopolisation) and female reproductive success.

In Model 4 (Table 1), our results indicated that the number of mating partners of the female’s dominant male had minimal effect on female reproductive success. This suggests that males that had more mating partners were not becoming sperm depleted or contributing to sperm limitation in females.

In Model 5 (Table 1), we found a negative interaction between the number of dam mates and skew in chick paternity on female chick production. This means that when females had a low number of mates, a higher skew in chick paternity positivity influenced female reproductive success, leading to increased chick production. However, as females engaged in more polyandrous mating behaviour, the positive impact of higher skew in chick paternity on female chick production decreased. Thus, the positive association between skew in chick paternity and female reproductive success weakened as females became more polyandrous (Figure 1).

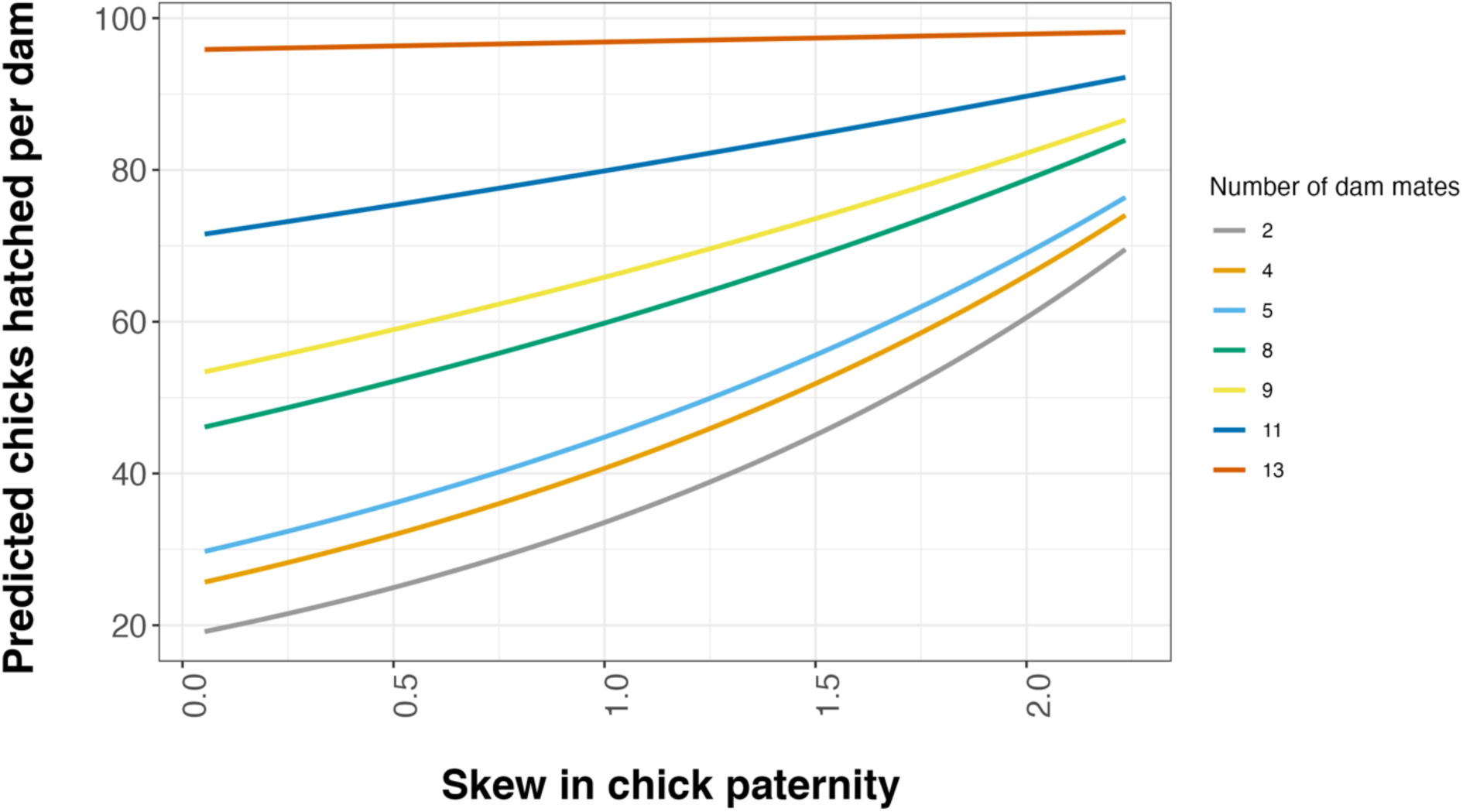

### Genetic variance in dam mating behaviours

For the total number of dam mating partners, we found genetic variance associated with Dam ID (Table 2), showing that the degree of female polyandry was partly influenced by genetic factors. We also observed substantial variance associated with Pen ID, suggesting that different patterns of social interactions may be occurring within different pens, contributing to differences in the amount of female polyandry observed among pens (Table 2).

**Table 2.**
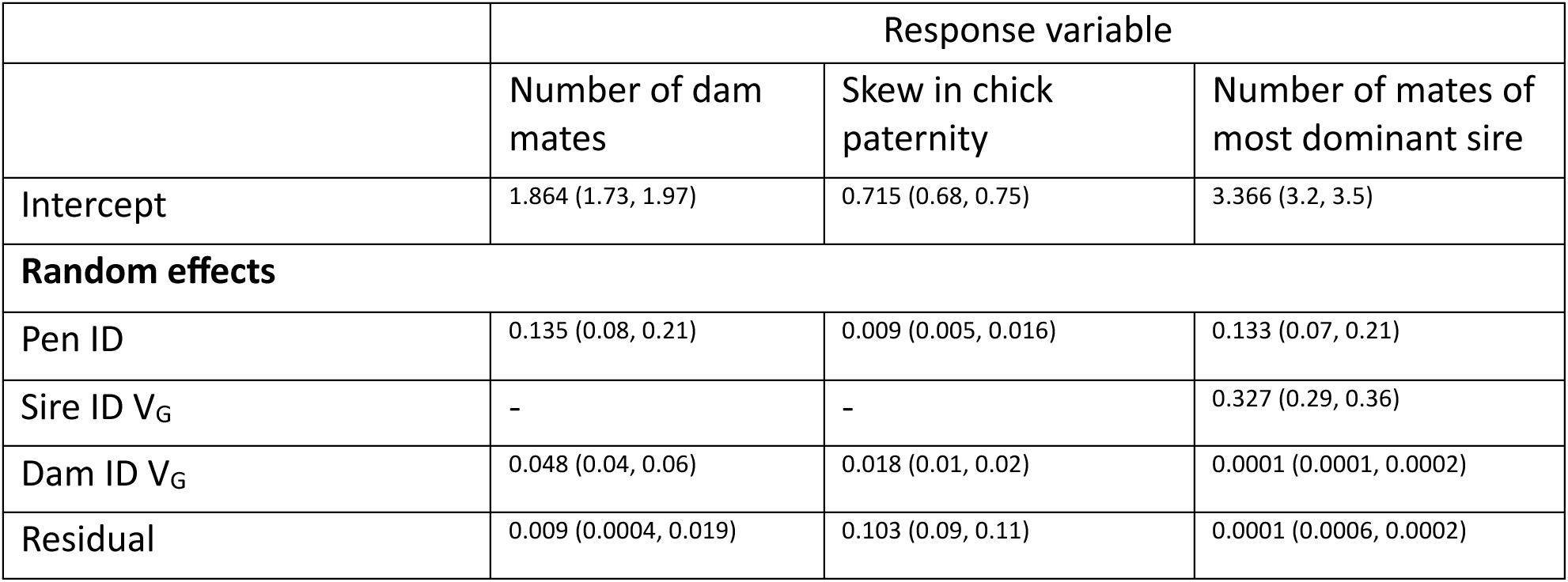
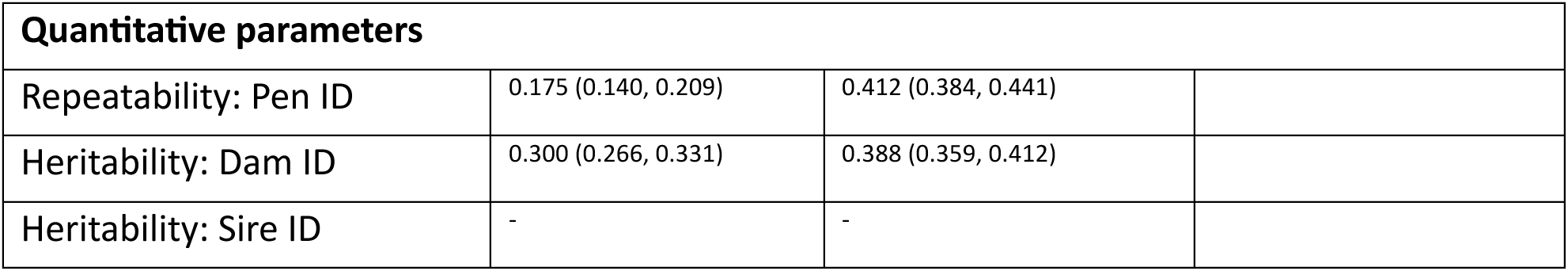
Results of three intercept-only animal models examining sources of genetic and environmental variance influencing number of dam mates, skew in chick paternity and number of mates of the most dominant sire. 95% credible intervals are reported in parentheses.

For skew in chick paternity, we found genetic variance associated with Dam ID, indicating that some females produce broods where the paternity of their chicks is more skewed towards a single male compared to other females (Table 2). This implies that genetic factors may influence the ability of females to have a larger proportion of their offspring sired from specific males. We also found environmental variance associated with Pen ID, indicating that social interactions within a pen may influence the degree of reproductive skew observed (Table 2). However, the genetic contributions of female identity towards reproductive skew were twice as large as those from Pen ID, suggesting that genetic factors, as opposed to social interactions within a specific pen, were more important in determining skew in chick paternity.

In terms of the number of mating partners of a female’s dominant male, we found strong genetic variance associated with Sire ID, suggesting that some males were genetically predisposed to attract and mate with multiple females (Table 2). We also observed a comparably low genetic variance associated with Dam ID, which might be expected, given that the number of sire mates is a predominantly male trait (Table 2). There was also some environmental variance associated with Pen ID, indicating that social interactions within specific pens may have contributed to the number of mating partners per sire (Table 2).

### Genetic covariance between mating behaviours in dams

The number of dam mates and skew in chick paternity were the two mating behaviours that impacted dam chick production the most. To understand how these behaviours phenotypically and genetically covaried with each other, we ran a bivariate model between these traits (Supplementary Materials, Table S1). We found a negative genetic covariance between these traits for DamID (-0.006, 95% CI: -0.008, -0.003). This negative genetic association indicates that higher levels of female polyandry may have come at the expense of reduced skew in chick paternity. We also observed a negative covariance between female polyandry and skew in chick paternity at the Pen ID level (-0.029, 95% CI: -0.05, -0.02). This indicates that the negative association between these two traits is not only genetic in origin but may also manifest because of differences in the types of social interactions that occurred within specific breeding pens.

### Mating behaviours influencing chick production in sires

Our findings support our initial sperm limitation prediction that higher rates of female polyandry are associated with an increase in female chick production. However, contrary to our second sperm limitation prediction, we found that higher skew in chick paternity was linked to an increase rather than a decrease in chick production. This suggests that factors beyond sperm limitation, such as genetic compatibility or cryptic female choice, may have contributed to the relationship demonstrated above between skew in chick paternity and female reproductive success in farmed Pekin ducks (see Discussion section).

To gain insight into how the number of mates and reproductive skew influenced male reproductive success, we examined the impact of mating behaviours on male reproductive success using five reciprocal models detailed in Table 3. Like female reproductive success, we found a positive effect of the total number of sire mating partners (Model 2, Table 3) and skew in chick maternity (Model 3, Table 3) on male reproductive success. In addition, we observed a negative interaction between the total number of sire mates and skew in chick maternity (Model 5, Table 3), indicating that the positive association between skew in chick maternity and male reproductive success weakened as males became more polygamous (i.e. a similar pattern to that shown in Figure 1 for females). We found that the average number of mating partners their females had tended to have limited influence on the number of chicks males sired (Model 4, Table 3). This indicates that males that were mating with more polygynous females were not suffering from reduced reproductive success because of potentially increased rates of sperm competition. Regarding the random effects, the base model (Model 1, Table 3) revealed genetic variance associated with Sire ID, indicating a genetic propensity for certain males to sire more chicks. Furthermore, there was environmental variance associated with Pen ID (Model 1, Table 3), suggesting that social interactions within pens may have contributed to the reproductive success of males across different pens.

**Table 3.** Results of five animal models of increasing complexity examining the impact of different mating behaviours (fixed effects) and sources of genetic and environmental variation (random effects) on sire chick production in farmed Pekin ducks. Model number corresponds to the model used in each analysis. 95% credible intervals are reported in parentheses.

| Parameter | 1 | 2 | 3 | 4 | 5 |
| --- | --- | --- | --- | --- | --- |
| **Fixed effects** |  |  |  |  |  |
| Intercept | 3.937 (3.77, 4.09) | 4.231 (4.13, 4.35) | 3.877 (3.76, 3.99) | 3.867 (3.76, 3.99) | 4.461 (4.31, 4.61) |
| Number of sire mates |  | 0.046 (0.04, 0.05) | 0.039 (0.03, 0.04) | 0.039 (0.03, 0.04) | 0.059 (0.05, 0.06) |
| Skew in chick maternity |  |  | 1.833 (1.08, 1.27) | 1.203 (1.09, 1.30) | 0.528 (0.36, 0.66) |
| Average number of mates of mating partners |  |  |  | 0.005 (-0.003, 0.013) | -0.001 (-0.009, 0.006) |
| Number of sire mates : Skew in chick maternity |  |  |  |  | -0.024 (-0.03, -0.02) |
| **Random effects** |  |  |  |  |  |
| Pen ID | 0.111 (0.04, 0.19) | 0.092 (0.05, 0.15) | 0.034 (0.01, 0.05) | 0.035 (0.02, 0.06) | 0.039 (0.02, 0.06) |
| Sire ID $V_G$ | 0.587 (0.46, 0.79) | 0.055 (0.03, 0.09) | 0.044 (0.02, 0.07) | 0.035 (0.02, 0.06) | 0.042 (0.03, 0.06) |
| Dam ID $V_G$ | 0.155 (0.05, 0.29) | 0.037 (0.03, 0.06) | 0.024 (0.01, 0.06) | 0.038 (0.03, 0.06) | 0.019 (0.008, 0.04) |
| Residual | 0.699 (0.48, 0.93) | 0.118 (0.07, 0.15) | 0.066 (0.03, 0.09) | 0.062 (0.03, 0.08) | 0.056 (0.03, 0.08) |

### Genetic variance in sire mating behaviours

We found that genetic variance associated with the number of sire mates was much larger than the environmental variance associated with Pen ID (Table 4). This indicates that certain males were genetically predisposed to attract and mate with a greater number of females, and that genetic sources of variance were more important than social interactions occurring within specific pens for this trait (Table 4). This differs somewhat from the variance pattern we observed in Dams (Table 2), where environmental variance associated with Pen ID was found to be larger than the genetic variance associated with Dam ID.

**Table 4.** Results of three intercept only animal models examining sources of genetic and environmental variance influencing number of sire mates, skew in chick maternity and average number of mates of mating partners. 95% credible intervals are reported in parentheses.

|  | Response variable |  |  |
| --- | --- | --- | --- |
|  | Number of sire mates | Skew in chick maternity | Average number of mates of mating partners |
| Intercept | 2.999 (2.85, 3.14) | 0.802 (0.76, 0.84) | 2.022 (1.92, 2.13) |
| PenID | 0.095 (0.04, 0.16) | 0.009 (0.003, 0.015) | 0.103 (0.05, 0.16) |
| SireID $V_G$ | 0.352 (0.23, 0.49) | 0.020 (0.01, 0.03) | 0.010 (0.004, 0.02) |
| DamID $V_G$ | - | - | 0.017 (0.01, 0.03) |
| Residual | 0.377 (0.27, 0.50) | 0.071 (0.06, 0.08) | 0.002 (0.0002, 0.004) |

Regarding skew in chick maternity, there was genetic variance associated with Sire ID, indicating that the maternity of some male’s chicks was more skewed towards a single dam compared to other males (Table 4). There was also environmental variance associated with Pen ID (Table 4). As with the results observed for females (Table 2), genetic variance was found to be more important than environmental sources of variance for this trait.

In terms of the average number of mates sire’s mating partners had, we found that environmental variance associated with Pen ID explained the largest source of variation for this trait, whilst a smaller proportion of genetic variance was explained by Dam ID and Sire ID (Table 4). This indicates that it is different social interactions within specific pens, as opposed to the genetic predisposition of males and females, which predominantly influences the average number of mating partners per female.

### Genetic covariance between mating behaviours in sires

Regarding the covariance between mating behaviours in sires (Supplementary Materials, Table S2), we found a negative genetic covariance between the number of sire mates and skew in chick maternity for Sire ID (-0.065, 95% CI: -0.08, -0.05). We also found a negative environmental covariance for same trait pair at the Pen ID level (-0.014, 95% CI: -0.03, - 0.005). These covariance results mirror those observed in dams (Supplementary Materials, Table S1), suggesting that the negative association between these traits is influenced by both genetic factors and differences in social interactions within specific breeding pens.

## Discussion

The main objective of this study was to assess the role of sperm limitation in contributing to the fertility gap in farmed Pekin ducks. In line with our initial prediction, we found a positive association between higher rates of polyandry and increased female reproductive success. This suggests that females use polyandry as a mating strategy to enhance their chances of acquiring sufficient sperm for fertilisation. However contrary to our expectations, we also found that female chick production increased with higher rates of skew in chick paternity, challenging the hypothesis that reduced reproductive success is due to limited sperm availability resulting from male monopolisation. Furthermore, we found no evidence that higher rates of male polygamy were associated with a decline in female chick production, implying that males mating with multiple females did not experience sperm depletion that contributed to sperm limitation in females. Collectively, our results highlight that that having multiple mating partners and some degree of reproductive skew was actually beneficial for chick production in female farmed Pekin ducks, suggesting that factors other than sperm limitation likely influence the size of any fertility gap in this system.

Our findings suggest that factors others than male monopolisation of females may have contributed to reproductive skew, as females gained fitness benefits when a higher proportion of their offspring were sired by a limited number of males. One potential factor that could explain this pattern is genetic compatibility (Kempenaers et al. 1999; Tregenza and Wedell 2000). Genetic compatibility refers to the likelihood of producing offspring when certain males and females mate and may arise due to recessive lethal alleles or intergenomic conflict leading to zygotes which are inviable or less viable (Zeh and Zeh 1996; Zeh and Zeh 1997; Kekäläinen 2021). Thus, it is possible that the increased skew in chick paternity and female reproductive success is the result of females mating with genetically compatible males (Kempenaers et al. 1999; Tregenza and Wedell 2000; Griffith et al. 2002). This may be because females benefit from increased heterozygosity in their broods as the proportion of deleterious alleles expressed is reduced (Kempenaers et al. 1999). Similar observations have been made in wild populations of adders, sand lizards and tree swallows where the proportion of a female’s viable offspring has been found to increase when females have a greater number of mating partners (Madson et al. 1992; Olsson et al. 1994; Kempenaers et al. 1999). Furthermore in tree swallows, fertilised unhatched eggs which failed to develop were found to be the result of embryo mortality, which declined with the number of mating partners a female had (Kempenaers et al. 1999).

Another possible explanation for the positive relationship between increased skew in chick paternity and female reproductive success is cryptic female choice (Springate and Frasier 2017). Cryptic female choice refers to the mechanism though which females can select and favour specific male sperm within their reproductive tract after copulation. This process could involve biasing or selecting for sperm from genetically compatible or superior males. Such biased sperm selection may increase the chances of producing viable offspring and hence increase female reproductive success.

In addition, male monopolisation of females may have fitness and individual welfare benefits. Mate guarding, as a form of male monopolisation, may serve to protect females from excessive male harassment or aggression(Gowaty and Buschhaus 1998). By preventing physiological stress in females, mat guarding could improve overall female condition and welfare, thereby further enhancing female reproductive success.

To test these alternative hypotheses then additional genetic, experimental and behavioural data would be required. In terms of genetic data, our current approach involves collecting DNA samples and performing parental assignment only on hatched chicks. However, it would be valuable to also collect DNA from fertilised eggs that fail to develop. This would identify instances where individuals have mated but failed to produce viable offspring and may shed light on potential genetic incompatibilities such as recessive lethal alleles. For experimental data, mating designs could be implemented where specific males and females are mated with each other,(e.g. the North Carolina 2 design) (Muthoni and Shimelis 2020). This approach would have the added benefit that matings could be controlled between particular male-female combinations, allowing us to determine which genotype combinations are the most and least compatible, or whether cryptic female choice is occurring in this system (Byrne et al. 2021).

Behavioural data could also provide valuable insight into the relationship between mating events and individual reproductive success. Utilising sensor technology would enable a high-throughput assessment of all mating events taking place in a pen and allow us to identify mating interactions between specific individuals that may provide cues to potential genetic compatibility between individual pairings (Toscano et al. 2019; van der Sluis et al. 2020; Carslake et al. 2022; Occhiuto et al. 2022). Furthermore, by examining the activity level and space use of individuals, we would be able to further identify whether certain movement-based behaviours are associated with more mating opportunities than others in both males and females (van der Sluis et al. 2020). Finally, by conducting more detailed behavioural observations on a subset of ducks, we could determine how specific types of mating interactions, such as male coercive or cooperative mating, impact female (and male) reproductive success, thus providing insights into cryptic female choice and its effect on the fertility gap in this system.

The presence of genetic variance in female polygamy and skew in chick paternity suggests that these traits could be selectively targeted in commercial breeding programmes to increase reproductive success in Pekin ducks. By focusing on individuals with genetic predispositions for high mate numbers or higher skew in chick paternity, breeders may be able to enhance chick production. However, our findings also highlight that social interaction in this case mean that it might not be so simple, and that female mate number is also influenced by factors that do not have a genetic basis, such as the social dynamics of the pen. Therefore, further investigation would be necessary into how the pen environment can be optimised (e.g., through different pen designs or social structures) to encourage high levels of female polygamy and hence chick production. This will enable breeders to optimise chick production by using a combined approach that looks to make genetic progress in relevant social behavioural traits, whilst providing a breeding environment that is conducive to effective mating behaviours.

It is also important to highlight here the negative covariance between female mate number and reproductive skew, which had a genetic and an environmental basis. From a behavioural perspective, this indicates that individuals may specialise in different breeding strategies, with some females mating with a high number of males, whilst other females may choose a small number of specific males to sire their majority of their offspring. From a commercial breeding perspective, this means that selection for genetic progress in one trait will come at the expense of the other, and so careful consideration needs to be given when implementing selection strategies for increased chick production. However, negative trait covariance does not mean that progress cannot be made in both traits, breeders need to phenotype both traits and select accordingly . Depending on the specific requirements of the breeding programme and the extent that these mating strategies also covary with other welfare and production related traits, emphasis may need to be placed upon promoting female polyandry or enhancing skew in chick paternity.

Building upon the findings from this study, future investigations could involve conducting individual-based data simulations to explore the performance of different reproductive strategies and their implications for commercial breeding programmes. By incorporating equations defining selecting gradients and trait covariance, along with statistical estimates obtained from duck data, this will provide greater insight into the potential benefits and trade-offs associated with different selection strategies. This approach will not only inform breeding programmes on how to enhance chick production but will also highlight the broader impact of specific breeding strategies on essential production traits and indicators of bird welfare.

## Conclusion

This study highlights factors influencing chick production and the fertility gap in farmed Pekin ducks. We show that both the number of dam mates and reproductive skew contribute to female reproductive success, indicating the potential for targeted selection strategies in commercial breeding programmes. However, our results challenge the sperm limitation hypothesis and the notion that male monopolisation negatively impacts female reproductive success. The exploration of alternative hypotheses such as genetic compatibility and cryptic female choice could help to explain these patterns, but this will require additional genetic, experimental and behavioural data. By integrating concepts from behavioural ecology, quantitative genetics and animal breeding, this study emphasises the importance of considering social behaviour in optimising commercial breeding programmes. Thus, this type of interdisplinary approach has the potential to open up new opportunities to develop effective breeding strategies that consider the genetic and (social) environmental factors impacting behavioural dynamics.

